# CD44 Controls Endothelial Proliferation and Functions as Endogenous Inhibitor of Angiogenesis

**DOI:** 10.1101/049494

**Authors:** Anne Pink, Marianna Školnaja, Taavi Päll, Andres Valkna

**Author notes:** Corresponding author: Andres Valkna.

## Abstract

CD44 transmembrane glycoprotein is involved in angiogenesis, but it is not clear whether CD44 functions as a pro- or antiangiogenic molecule. Here, we assess the role of CD44 in angiogenesis and endothelial proliferation by using *Cd44*-null mice and CD44 silencing in human endothelial cells. We demonstrate that angiogenesis is increased in *Cd44*-null mice compared to either wild-type or heterozygous animals. Silencing of CD44 expression in cultured endothelial cells results in their augmented proliferation and viability. The growth-suppressive effect of CD44 is mediated by its extracellular domain and is independent of its hyaluronan binding function. CD44-mediated effect on cell proliferation is independent of specific angiogenic growth factor stimulation. These results show that CD44 expression on endothelial cells constrains endothelial cell proliferation and angiogenesis. Thus, endothelial CD44 might serve as a therapeutic target both in the treatment of cardiovascular diseases, where endothelial protection is desired, as well as in cancer treatment, due to its antiangiogenic properties.

## Introduction

Angiogenesis is a pathophysiological process involved in wound healing as well as tumor growth and metastasis. The switch of normally quiescent blood vessels to angiogenesis is determined by the balance of proangiogenic and antiangiogenic factors in tissue microenvironment. In tumors, endothelial cell (EC) proliferation and survival is regulated by growth factors such as VEGF, FGF-2, or HGF, released either by tumor cells or normal cells in the tumor stroma. A survival benefit has been shown for blocking proangiogenic signaling by VEGF inhibitors in tumor therapy and continuation of such therapy beyond progression [1]. However, tumors can be intrinsically resistant to anti-VEGF therapy or acquire resistance through different mechanisms during the therapy [2]. Better understanding of microenvironmental or EC-specific factors involved in the regulation of EC proliferation and blood vessel formation is needed to develop new more effective antiangiogenic tumor therapies.

CD44 cell-surface glycoprotein has been shown to be necessary for efficient tumor vascularization [3] and to mediate induction of angiogenesis in response to hyaluronan (HA) oligomers [4]. CD44 mediates cell adhesion to its principal ligand HA via its N-terminal HA-binding domain (HABD) [5, 6]. CD44 and HA interaction is important in the immune response where it mediates leukocyte rolling on HA during leukocyte recruitment into the inflammatory site [7–9]. CD44 also mediates HA-induced effects on vasculature. *In vivo* silencing of endothelial CD44 resulted in reduced vascular density of Matrigel plug implants in response to low molecular weight HA [4]. The interaction of CD44 with high molecular weight HA inhibits vascular smooth muscle cell proliferation, and CD44 deficiency has been shown to increase neointima formation after arterial injury [10]. *Cd44*-null mice display reduced vascularization of Matrigel plugs containing CD44-positive B16 melanoma cells, as well as reduced vessel density in B16 melanoma and ID8-VEGF ovarian carcinoma xenografts [3]. However, the contribution of CD44 to *in vivo* angiogenesis has not been studied in *Cd44*-null mice in less complex models using defined angiogenic growth factors instead of large numbers of tumor cells.

We have previously shown that administration of recombinant soluble CD44 hyaluronan binding domain (HABD) inhibits *in vivo* angiogenesis in both VEGF- and FGF2-stimulated chick chorioallantoic membranes and tumor xenograft growth, and that *in vitro*, sCD44 HABD controls EC proliferation [11]. Significant levels of soluble CD44 have been reported in normal human serum and even higher levels in mouse serum. Increased serum levels of soluble CD44 (sCD44) have been linked to different pathological conditions, including cancer and type 2 diabetes [12–14]. Soluble CD44 is mainly a shedding product of membrane CD44 [15, 16]. If the hypothesis of the proangiogenic role of CD44 holds, recombinant soluble CD44 HABD should function as a decoy receptor for HA, the principal ligand of CD44. Nevertheless, we have previously found that the non-HA binding mutant of CD44 HABD showed similar antiangiogenic and tumor growth inhibitory functions [11], suggesting that a mechanism other than HA binding might be involved.

The aim of the current study was to analyze the effects of *Cd44* gene deficiency and non-HA-binding CD44 HABD treatment on *in vivo* angiogenesis and *in vitro* EC proliferation to conclusively determine the role of CD44 in angiogenesis and in order to cast light upon the mode of action of the non-HA-binding CD44 HABD.

## Results

### *Cd44*-null mice display increased angiogenic response

We studied angiogenesis in mice lacking CD44 (*Cd44*^−/−^). In a preliminary experiment, we assessed the invasion of isolectin-B4- and CD105-positive cells into the Matrigel plug in response to 50 ng/ml VEGF. Surprisingly, results indicated that plugs from *Cd44*^−/−^ mice contained more cells than plugs from wild-type mice (data not shown). We also tested capillary outgrowth in aortic fragments isolated from *Cd44*^−/−^ mice. Aortic fragment assay indicated apparently more robust outgrowth of capillary-like structures from *CD44*^−/−^ fragments compared to wild-types (Supplemental Fig. 1 in Online Resource 1). This finding is in agreement with Chun et al. 2004 study [17], showing no defect in capillary outgrowth and vessel morphogenesis in aortic fragments isolated from *Cd44*^−/−^ mice. In the following series of experiments, we used the directed *in vivo* angiogenesis assay [18] to quantitatively test FGF-2/VEGF-induced angiogenic response in *Cd44*^−/−^ and wild-type mice of C57BL/6, C3H or mixed genetic backgrounds (Fig. 1). We found a 5.4 ± 2.2-fold increase in blood vessel invasion in response to FGF-2/VEGF compared to unstimulated controls in *Cd44*^−/−^ mice of mixed genetic background (t-test: *P* = 0.048, N = 2 experiments, effect size 1.2 ± 0.45 [95% CI, 0.37-2.1]; Fig. 1A). Angiogenesis induction in wild-type mouse strains compared to unstimulated controls was in C3H 1.4 ± 0.5-fold (t-test: *P* = 0.41, effect size 0.081 ± 0.071 [95% CI, −0.061-0.22]), C57BL/6 1.5 ± 0.62-fold (*P* = 0.38, effect size 0.2 ± 0.17 [95% CI, −0.09-0.56]), and in wild-type mice of mixed genetic background 4.2 ± 1-fold (t-test: *P* = 0.13, effect size 0.14 ± 0.039 [95% CI, 0.082-0.2]; Fig. 1A). The estimated effect size of growth factor stimulation was large in *Cd44*^−/−^ mice with confidence interval ranging from small to very large, whereas in wild-type mouse strains growth factor stimulation displayed small effect size with confidence intervals including zero. When comparing growth factor-stimulated groups, *Cd44*^−/−^ mice displayed apparently severalfold higher blood vessel invasion into angioreactors in response to FGF-2/VEGF than wild-type mice. These data suggest that *Cd44* deficiency results in augmented angiogenic response. However, factors other than CD44 might also affect the efficiency of angiogenesis induction in mice of different genetic backgrounds [19].

**Fig. 1.**
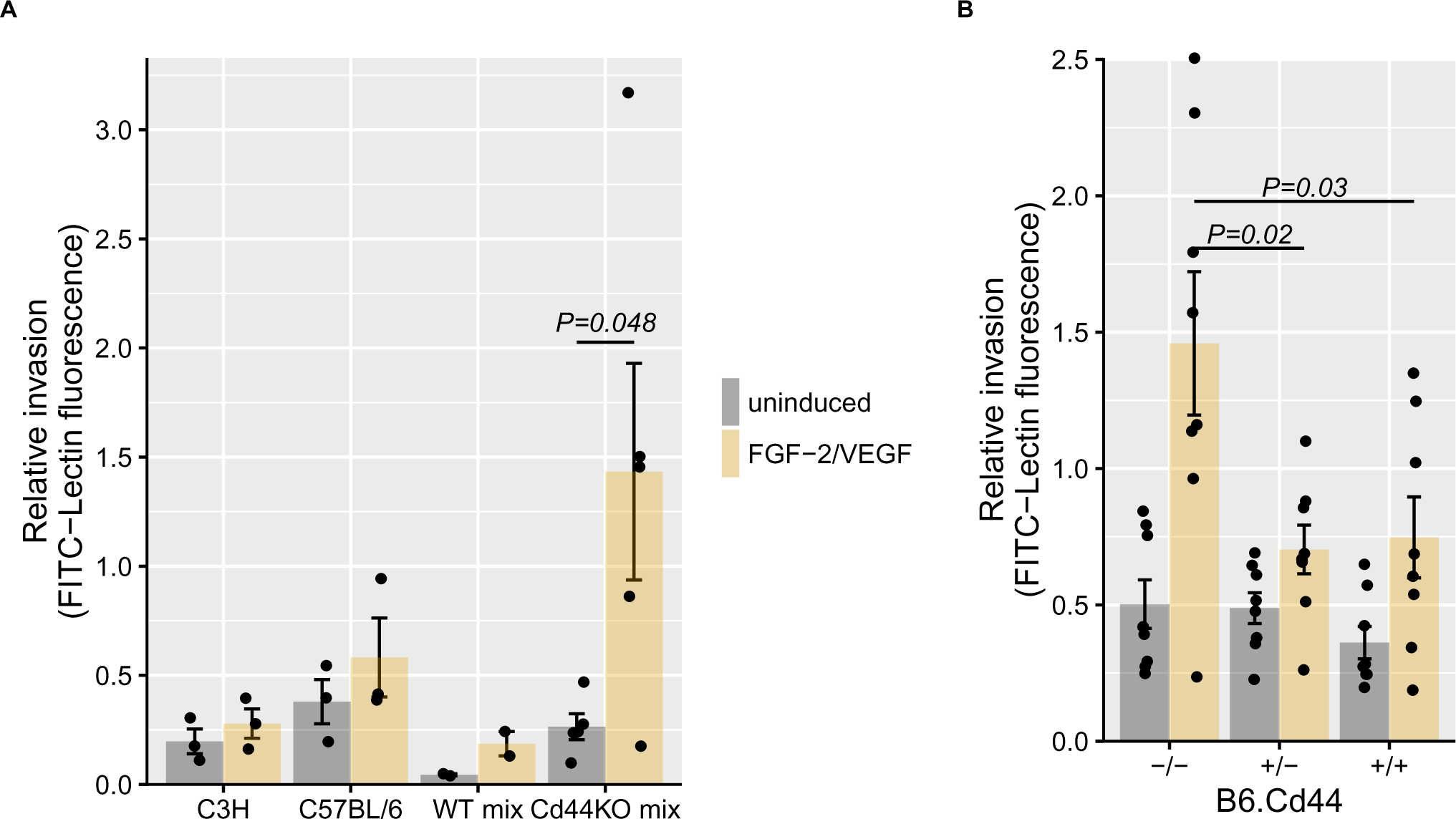
Increased angiogenesis in CD44-null mice. (**A**) Angiogenesis was analysed in C3H, C57BL/6, and in wild-type and *Cd44*^−/−^ mice of mixed genetic backgrounds (WT mix and Cd44KO mix, respectively). Basement membrane extract–filled angioreactors containing premixed FGF-2, VEGF and heparin or PBS for uninduced controls were implanted SC into the flanks of mice. Each mouse received 2 angioreactors, 1 per flank. 14 days after implantation, angioreactors were resected and the population of ECs within the angioreactor matrix was assessed by FITC-lectin staining. The number of fluorescent cells was quantitated by microplate reader. Raw readings from independent experiments were scaled by dividing by their quadratic mean. N = 2–5 mice per condition from 2 independent experiments. P value is from Student’s t-test. (**B**) Angiogenesis in *Cd44*^−/−^ mice and their heterozygous (*Cd44*^+/−^) and wild-type (*Cd44*^+/+^) littermate that had been backcrossed six generations to the C57BL/6 background. The data are represented as the mean ± SEM. Each dot represents the mean of two angioreactors for an individual mouse. N = 8 mice per condition from 2 independent experiments. P values are from ANOVA post hoc comparisons using the Tukey HSD test.

To confirm increased angiogenesis in the absence of CD44, and to study the contribution of CD44 to neoangiogenesis in an inbred, genetically homogeneous background, we backcrossed *Cd44*^−/−^ mice for six generations to the C57BL/6 strain. Using littermate controls, we found that *Cd44* genotype had a significant effect on angiogenesis induction (ANOVA, F_2,42_ = 3.98, *P* = 0.026). Angiogenesis was increased in *Cd44*-null animals compared to wild-type or heterozygous mice. FGF-2/VEGF stimulation resulted in a 2.1 ± 0.5-fold increase of blood vessel invasion in C57BL/6 *Cd44*^+/+^ mice (one-way ANOVA, *P* = 0.03; effect size 0.39 ± 0.15 [95% CI, 0.098-0.67]); in a 1.4 ± 0.24-fold increase in their *Cd44*^+/−^ littermates (one-way ANOVA, *P* = 0.061; effect size 0.22 ± 0.1 [95% CI, 0.019-0.4]); and in a 2.9 ± 0.75-fold increase in their *Cd44*^−/−^ littermates (one-way ANOVA, *P* = 0.0039; effect size 0.96 ± 0.26 [95% CI, 0.45-1.5]); N = 8 mice per genotype from 2 independent experiments. Angiogenesis induction was associated with *Cd44* genotype in the growth factor-induced group of mice (ANOVA, F_2,21_ = 5.45, *P* = 0.012), whereas there was no difference in baseline vessel invasion between different *Cd44* genotypes in the unstimulated group. Post-hoc testing of pairwise differences indicated that *Cd44*^−/−^ mice displayed increased angiogenesis compared to either their *Cd44*^+/−^ or *Cd44*^+/+^ littermates (Fig. 1B).

We conclude from these experiments that CD44 deficiency results in augmented angiogenesis induction; and thus CD44 functions as an endogenous angiogenesis inhibitor.

### Recombinant CD44 non-HA-binding mutant Fc-fusion protein inhibits angiogenesis *in vivo*

We studied whether increasing the dose of CD44 by systemic administration of its soluble analog could suppress angiogenesis. We have previously shown that bacterially expressed GST-tagged non-HA-binding CD44 (CD44-3MUT) inhibits angiogenesis in chick chorioallantoic membrane and subcutaneous tumor xenograft growth in mice [11]. Untagged and GST-tagged CD44-3MUT display very short serum half-life limiting their potential *in vivo* use [20]. Thus, in order to improve *in vivo* efficacy, we generated CD44-3MUT with a C-terminal fusion of human IgG1 Fc region (CD44-3MUT-Fc). Regarding pharmacokinetic characterization of CD44-3MUT-Fc, its serum half-life after intravenous administration to rats was 17 min (Supplemental Fig. 2 in Online Resource 1). The volume of distribution of CD44-3MUT-Fc was 18% of total body weight (%TBW), suggesting improved biodistribution compared to untagged CD44-3MUT protein (1.8% TBW; [20]).

We tested the antiangiogenic effects of CD44-3MUT-Fc in athymic nude mice. The results showed that FGF-2/VEGF stimulation led to a 4.5 ± 1.1-fold increase of blood vessel invasion into angioreactors over the PBS treated control groups (t-test: *P* = 7.7e-07; effect size 0.8 ± 0.15 [95% CI, 0.5-1.1]; Fig. 2B, C). We observed that intraperitoneal treatment of mice with CD44-3MUT-Fc inhibited FGF-2/VEGF-induced angiogenesis. In the treatment group receiving 25 mg/kg CD44-3MUT-Fc, angiogenesis was inhibited to the unstimulated basal level compared to the growth factor-induced PBS treatment group (t-test: *P* = 1.6e-05; effect size −0.8 ± 0.16 [95% CI, −1.1–0.5]). Administration of 0.5 or 5 mg/kg CD44-3MUT-Fc also resulted in apparent angiogenesis inhibition, but the response was less robust (Fig. 2C). For recombinant protein control treatments, we used irrelevant rhIgG mAb or rhIgG-Fc. Both these molecules were purified identically to CD44-3MUT-Fc. The 0.5 mg/kg CD44-3MUT-Fc treatment showed significant inhibitory effect compared to pooled 0.5 mg/kg rhIgG/rhIgG-Fc control treatment group (t-test: *P* = 0.034; effect size 0.5 ± 0.24 [95% CI, 0-0.9]). Mice receiving intraperitoneally 5 or 15 mg/kg doses of rhIgG-Fc showed similar angiogenic response as PBS and 0.5 mg/kg rhIgG-Fc control treatment groups, suggesting that the IgG-Fc portion of CD44-3MUT-Fc is not responsible for the effects observed.

**Fig. 2.**
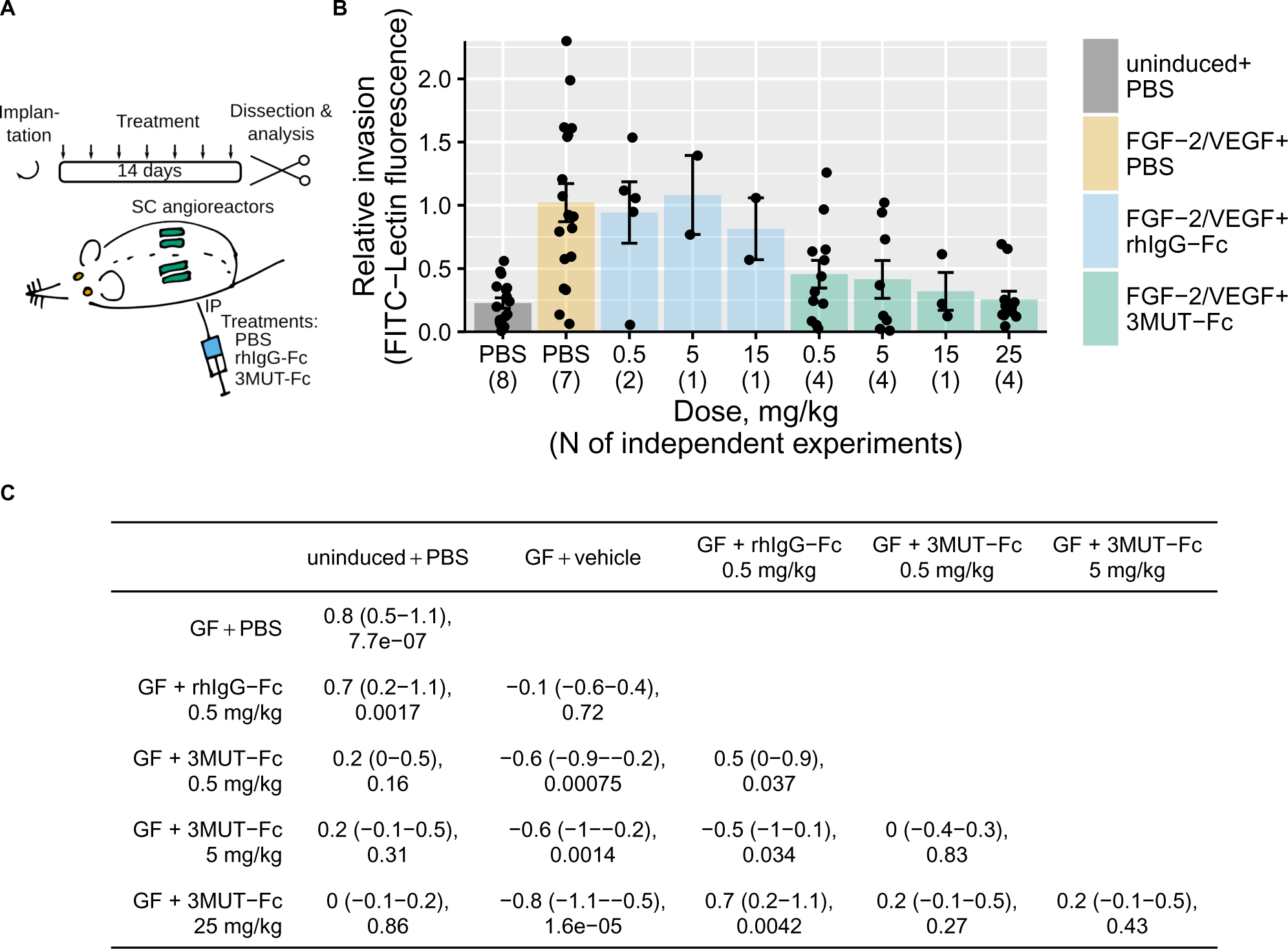
Recombinant CD44-3MUT Fc fusion protein inhibits angiogenesis *in vivo*. First, the assay and quantitation were performed similarly to those described in Fig. 1, except that each mouse received 4 angioreactors, 2 per flank. The day after implantation, the mice started to receive CD44-3MUT-Fc, control (rhIgG1-Fc) or vehicle (PBS) every second day via IP injections for 14 days. (**A**) Schematic presentation of experimental design. (**B**) Relative blood vessel invasion into matrix–filled angioreactors. The data are represented as the mean ± SEM. Datapoints show the mean of 4 angioreactors for individual mice. N – the number of independent experiments. GF – growth factors (FGF-2/VEGF). (**C**) Effect size with 95% confidence intervals (upper row) and P values (lower row) of pairwise comparisons of the data shown in (**B**). Effect sizes were calculated by Cohen’s d formula. Confidence intervals were derived using bootstrap resampling. P values are from t tests using pooled SD.

Together, our results show that systemic administration of CD44-3MUT-Fc effectively inhibits *in vivo* angiogenesis.

### Soluble CD44 levels are not affected by angiogenesis

Serum levels of sCD44 are increased due to enhanced CD44 shedding in case of inflammation or tumor growth. Previous research suggests that serum sCD44 concentrations show substantial variability between different mouse strains, and are significantly reduced (< 1 μg/ml) in severely immunodeficient mice [21]. To assess the significance of sCD44 in angiogenic response, we determined sCD44 concentrations in the sera of different wild-type mouse strains, in *Cd44*^+/−^ mice, and in athymic nude mice. Sera from *Cd44*^−/−^ mice were used as negative controls. We found that serum concentrations of sCD44 were similar in athymic nude mice and in wild-type mouse strains (Fig. 3A and Table 1). Soluble CD44 levels in *Cd44*^+/−^ animals were reduced on average by 35% (95% credible interval: 16-53%) compared to *Cd44*^+/+^ mice.

**Fig. 3.**
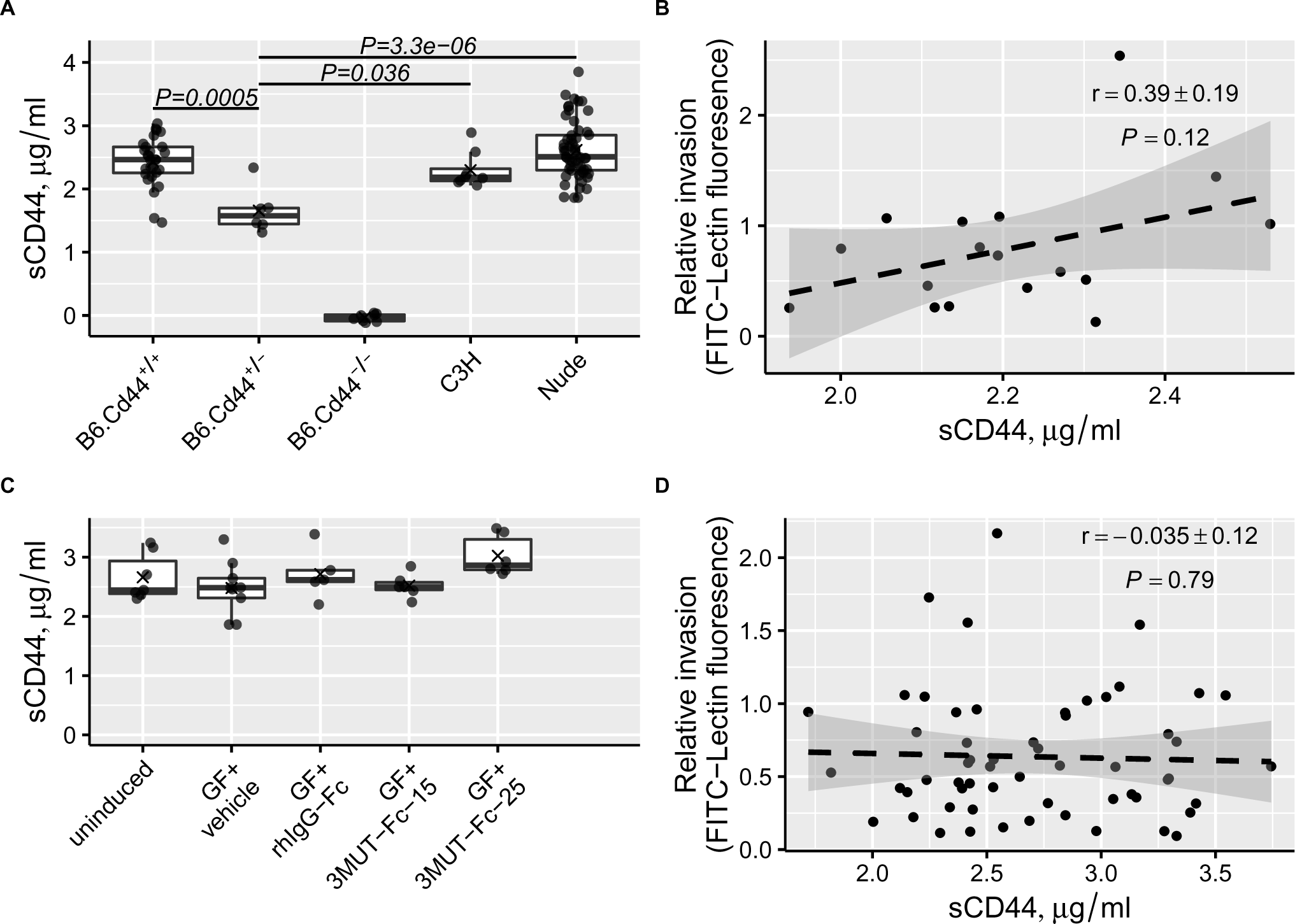
Soluble CD44 concentrations in mouse serum. Blood was collected from mice of different genetic backgrounds and the mice used in the angiogenesis experiments shown in Figs 1 and 2. (**A**) Serum levels of soluble CD44 in mice from different strains. Each dot represents an individual mouse. Cross indicates the mean. P values are from ANOVA post hoc comparisons using the Tukey HSD test. (**B**) The correlation between relative blood vessel invasion and post-experiment serum sCD44 in wild-type mice. *Cd44*-null mice and nude mice were excluded from the dataset. Pearson’s r and the associated P values are shown. (**C**) Post-experiment serum levels of sCD44 in nude mice from different treatment groups. Treatments where more than five mice were analysed are shown. Each dot represents an individual mouse. Cross indicates the mean. GF – growth factors (FGF-2/VEGF). (**D**) The correlation between relative blood vessel invasion and post-experiment serum sCD44 in nude mice. Pearson’s r and the associated P values are shown. In (**B** and **D**), dashed line is the linear model fit, gray shading is the standard error interval of fitted values.

**Table 1.** Soluble CD44 level in mouse serum. Concentration is shown as mean ± SEM.

| Strain | sCD44, $\mu\text{g/ml}$ | N | 95% credible interval |
| --- | --- | --- | --- |
| B6 | 2.4 ± 0.07 | 28 | 2.3-2.6 |
| B6.Cd44<sup>+/-</sup> | 1.7 ± 0.1 | 6 | 1.2-2.1 |
| C3H | 2.3 ± 0.1 | 8 | 2-2.6 |
| Nude | 2.6 ± 0.06 | 55 | 2.5-2.7 |

To analyse whether sCD44 concentrations correlate with angiogenesis induction, we excluded *Cd44*^−/−^ mice and nude mice from the dataset. We found no correlation between relative blood vessel invasion and post-experiment serum levels of sCD44 (Fig. 3B). Next, we evaluated whether the induction of angiogenesis, rhIgG-Fc or systemic treatments with CD44-3MUT-Fc that we used in the *in vivo* angiogenesis model lead to changes in the serum levels of sCD44 in nude mice. This analysis showed that neither angiogenesis induction nor treatments had any effect on the serum concentrations of sCD44 in nude mice (ANOVA, F_11,32_ = 0.975, *P* = 0.49; Fig. 3C). It also revealed that there was no correlation between post-experiment serum concentrations of sCD44 and vessel invasion into angioreactors irrespective of experimental intervention (Fig. 3D).

### CD44-3MUT-Fc inhibits endothelial cell proliferation and viability

To find out whether CD44-3MUT-Fc-mediated inhibition of angiogenesis is caused by its effects on ECs, we tested CD44-3MUT-Fc in a cell proliferation assay. We used cells synchronized by serum starvation to model the initial stages of stimulation of quiescent endothelial cells (Fig. 4A). We applied different concentrations of CD44-3MUT-Fc to growth arrested HUVECs and released cells from arrest by stimulation with 25 ng/ml VEGF. Realtime growth curves of untreated controls show that 25 ng/ml VEGF induces robust proliferation in HUVECs that is sustained for at least 72 h (Fig. 4B, leftmost panel). In contrast, CD44-3MUT-Fc treatment dose-dependently suppressed 25 ng/ml VEGF-stimulated HUVEC growth, compared to rhIgG-Fc control treatments (Fig. 4B). The difference in growth kinetics between CD44-3MUT-Fc and rhIgG-Fc treatments became apparent approximately 24 h after VEGF-induced release of cells from arrest. After this timepoint, rhIgG-Fc control-treated cells continued to proliferate, but in CD44-3MUT-Fc-treated wells cell density plateaued. Next, we used the same growth arrested HUVEC model in a cell proliferation and viability assay to compare CD44-3MUT-Fc efficacy in inhibiting cell proliferation stimulated by either FGF-2, VEGF or HGF (Fig. 4C to E). We used an endothelial-specific inhibitor of cell proliferation, fumagillin, as a positive control to define the maximum response in our assay. FGF-2 and VEGF induced robust proliferation in growth-arrested HUVECs, whereas HGF-stimulation resulted in much lower cell proliferation (Fig. 4C to E, left panels). Compared to rhIgG-Fc, CD44-3MUT-Fc dose-dependently inhibited HGF-stimulated proliferation with the maximum inhibition of 60% ± 12.6 after 72 h incubation. FGF-2 or VEGF-stimulated EC growth was inhibited less efficiently by CD44-3MUT-Fc when compared to rhIgG-Fc, as the growth was reduced by a maximum of 10.7% ± 2.4 and 13.9% ± 4.5, respectively (Fig. 4B to E, and G). These results are in agreement with respective growth factor potencies to stimulate HUVEC proliferation, CD44-3MUT-Fc was less efficient in inhibiting FGF-2- or VEGF-induced proliferation, and showed more efficacy in case of a weak inducer, HGF.

**Fig. 4.**
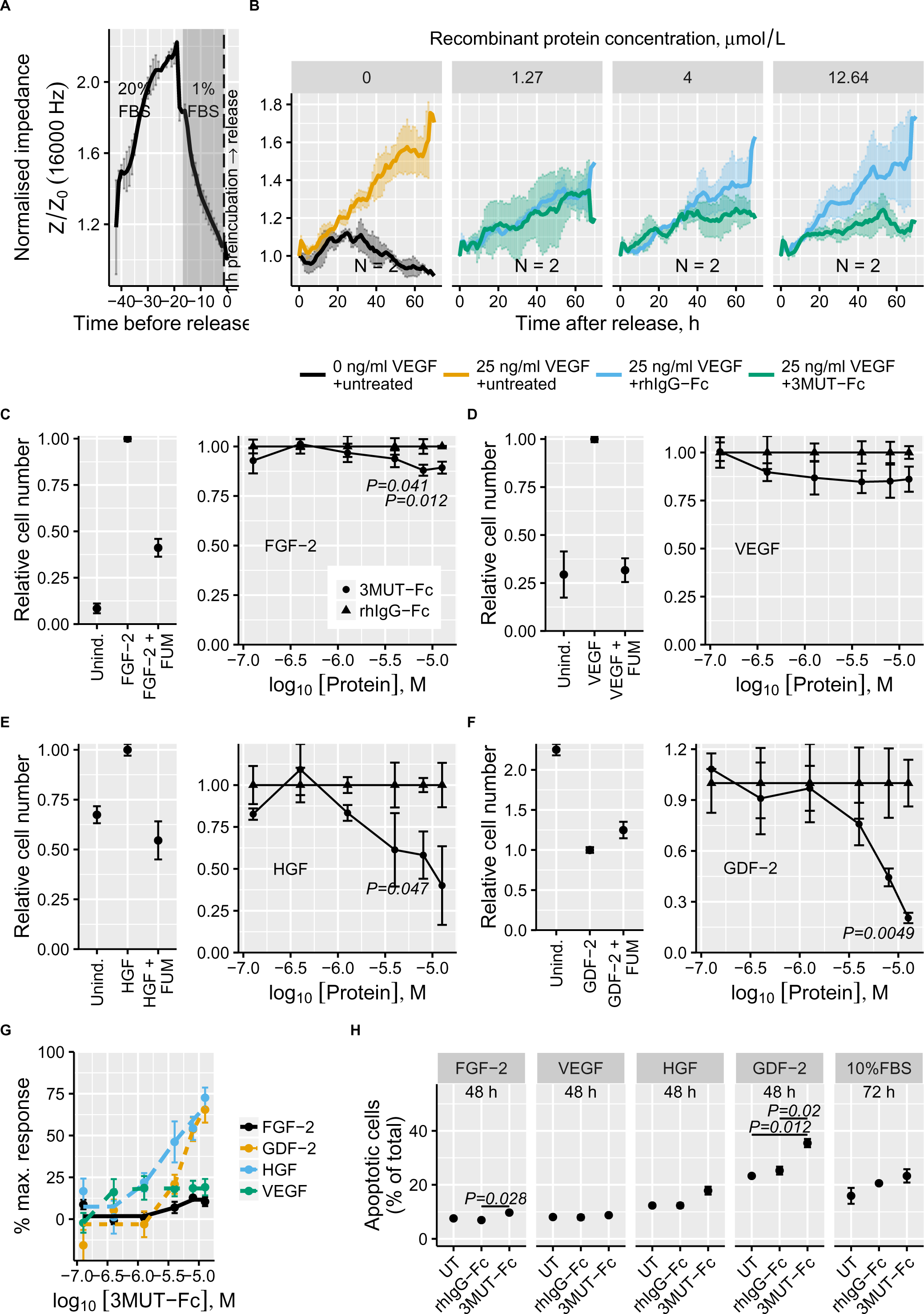
CD44-3MUT-Fc inhibits EC growth. (**A**) Real-time track of cell adhesion and synchronisation of HUVECs seeded onto 96-well electrode arrays. After seeding, the cells were grown for about 24 h. After that the cells were starved overnight in the media supplemented with 1% FBS (gray area). 1 h before the release from serum starvation (vertical dashed line) the cells were preincubated with different concentrations of rhIgG-Fc or CD44-3MUT-Fc in 5% FBS–containing media. (**B**) Growth curves of HUVECs released from serum starvation by supplementing preincubation media with 25 ng/ml VEGF. Facet labels show rhIgG-Fc or CD44-3MUT-Fc concentrations during preincubation. The data are represented as the mean ± SEM. N – the number of independent experiments. (**C–F**) HUVECs were synchronised and pretreated as in panel A. After preincubation, the cells were stimulated either with 25 ng/ml FGF-2 (**C**), 25 ng/ml VEGF (**D**), 63 ng/ml HGF (**E**) or 10 ng/ml GDF-2 (**F**). After 72 h, the number of viable cells was quantitated by measuring the ATP per well. Left: the effect of growth factor stimulation and the effect of 10 nM fumagillin (FUM) as a positive control for inhibition of cell proliferation. Right: the dose-response curves of rhIgG-Fc (filled triangles) and CD44-3MUT-Fc (filled circles). The data are represented as the mean ± SEM. N = 3–4 independent experiments. (**G**) CD44-3MUT-Fc dose-response curves for FGF-2, GDF-2, HGF or VEGF stimulated HUVEC. The data are represented as the mean ± SEM. (**H**) Apoptosis of HUVECs stimulated with different growth factors and treated with 12.64 μM (-4.9 log_10_ M) rhIgG-Fc, CD44-3MUT-Fc or left untreated. Apoptosis was quantitated by Annexin V staining. The data are represented as the mean ± SEM. N = 2 independent experiments. P values are from the ANOVA post hoc comparisons using the Tukey HSD test.

CD44v6 interacts with VEGFR2 and MET [22]. Therefore, we tested whether CD44-3MUT-Fc has any effect on the protein levels or activation of these receptors. Western blot analysis showed no change in either VEGFR2 or MET protein levels or receptor activation in response to CD44-3MUT-Fc (Supplemental Fig. 3 in Online Resource 1). This suggests that CD44-3MUT-Fc does not inhibit EC growth by direct targeting of growth factor receptor signaling pathways. Additionally, we tested the effect of CD44-3MUT-Fc on GDF-2-stimulated HUVECs (Fig. 4D). Vascular quiescence factor GDF-2 (BMP-9) belongs to the TGF-β superfamily ligands and regulates angiogenesis via ALK1, a type 1 TGF-β receptor [23, 24]. As shown in the left panel of Fig. 4F, in our model GDF-2 is strongly anti-mitotic and induces the cell cycle block. Compared to rhIgG-Fc control treatments, CD44-3MUT-Fc showed a dose-dependent inhibitory effect on cell numbers in GDF-2-treated HUVECs with 79.6% ± 6.7 of maximum response (Fig. 4F and G).

To ascertain whether apoptosis contributes to CD44-3MUT-Fc-induced growth inhibition, we used Annexin V-FITC staining. We found that upon release from serum starvation, the basal levels of apoptosis in the HUVEC population were inversely related to the growth factor potency to stimulate cell proliferation. In response to incubation with 12.64 μM (-4.9 log_10_ M) CD44-3MUT-Fc, the number of apoptotic cells relative to pooled control treatments was increased by 9% ± 5 in VEGF, 28% ± 14 in 10%FBS, 34% ± 6 in FGF-2, 45% ± 12 in HGF, 46% ± 6 in GDF-2-stimulated cells. However, VEGF- or FGF-2-induced cells were the most protected against apoptosis induced by CD44-3MUT-Fc-treatment (Fig. 4H). In contrast, GDF-2-mediated growth arrest enforced cells to undergo apoptosis and this trend was further increased by CD44-3MUT-Fc treatment. The observed increase in apoptosis from a relatively low basal level in response to CD44-3MUT-Fc treatment and its apparent correlation with growth factor potency to stimulate proliferation, suggest that apoptosis occurs secondary to CD44-3MUT-Fc-mediated inhibition of cell proliferation.

Collectively, our data show that CD44-3MUT-Fc inhibits EC proliferation.

### CD44 is not involved in GDF-2/ALK1-dependent SMAD signaling

Several studies suggest that CD44 is associated with TGF-β signaling, since the cytoplasmic tail of CD44 directly interacts with SMAD1 [25], CD44 forms a galectin-9-mediated complex with BMPR2 [26], and HA induces CD44 to complex with TGFBR1 [27]. Given that CD44-3MUT-Fc treatment resulted in an enhanced growth inhibitory effect in GDF-2-arrested HUVECs, we wanted to test whether CD44 could be involved in GDF-2 mediated SMAD activation. We studied pSMAD1/5 nuclear localization and SMAD1/5 target gene activation in response to GDF-2 stimulation in CD44-silenced HUVECs. CD44-targeting siRNA (siCD44) transfection resulted in substantial CD44 protein or mRNA downregulation compared to non-targeting siRNA control (siNTP) (Supplemental Fig. 4 in Online Resource 1). Simultaneously, we detected a robust GDF-2-dependent pSMAD1/5 nuclear localization (Supplemental Fig. 4A and B in Online Resource 1) and an induction or repression of selected known SMAD1/5 target genes ID1, SMAD6, SMAD7 or c-MYC, respectively (Supplemental Fig. 4C in Online Resource 1). Immunofluorescence analysis showed that the nuclear area or other size/shape parameters of cell nuclei did not differ between siCD44 or siNTP control-silenced cells (data not shown). We found that silencing of CD44 did not affect the nuclear localization of pSMAD1/5 or SMAD1/5 target gene expression in response to GDF-2 stimulation (Supplemental Fig. 4A to C in Online Resource 1).

Next, we studied the effect of CD44-3MUT-Fc on SMAD1/5 signaling by using a BMP-responsive element reporter (BRE). We found that BRE reporter activity in HUVECs was increased in response to GDF-2 stimulation, but this response was not sensitive to either CD44-silencing or CD44-3MUT-Fc treatment (Supplemental Fig. 5A in Online Resource 1). In line with this, Western blot analysis of GDF-2 stimulated HUVECs treated with CD44-3MUT-Fc showed no change in pSMAD1/5 levels (Supplemental Fig. 5B in Online Resource 1). To test whether prolonged CD44-3MUT-Fc exposure *in vivo* could trigger changes in SMAD1/5-mediated gene expression, we analysed lung tissue of nude mice from two angiogenesis experiments described in Fig. 2 for the expression of selected SMAD target genes. We found no changes in the expression levels of SMAD1/5 or NF-κB target genes (Supplemental Fig. 5C in Online Resource 1).

Together, these results suggest that neither CD44 nor CD44-3MUT-Fc are involved in GDF-2-ALK1-SMAD1/5 signaling in ECs.

### CD44 silencing augments endothelial cell proliferation

Since angiogenesis assay results showed increased blood vessel invasion in *Cd44*^−/−^ mice and inhibition of this response by CD44-3MUT-Fc treatment, we wanted to test whether CD44 knockdown in ECs results in increased cell growth. To this end, we used siRNA transfected HUVECs that had been growth arrested by serum starvation. Growth-arrested cells were released by the addition of either 20% FBS or different concentrations of VEGF or FGF-2. Real-time impedance measurements showed that compared to 5% FBS stimulation (Fig. 5A, left), 20% FBS or growth factor supplementation released cells from the cell-cycle block and stimulated their growth sustainably over 72 h (Fig. 5A and C). We found that siCD44 transfected cells reached higher densities at 72 h than control siNTP-transfected cells. Augmented cell growth and higher cell density in siCD44-transfected ECs at the end of the experiment was independent of type and concentration of the growth factor used for stimulation (Fig. 5A, C and E). End-point quantitation of viable cells performed 72 h after release supported the impedance measurement results. siCD44-transfected HUVECs displayed increased cell numbers over all tested growth factors and concentrations (Fig. 5B and D). These data suggest that the effect of CD44 silencing on cell proliferation was additive to the stimulatory effect of growth factors. The additive effect of CD44 silencing on cell proliferation was further supported by the typical VEGF dose-dependent flattening of the growth curve that was observed in case of VEGF-stimulated HUVECs. In case of FGF-2, such suppression did not occur, and cell density increased linearly with growth factor concentration within the range (8 to 79 ng/ml) tested.

**Fig. 5.**
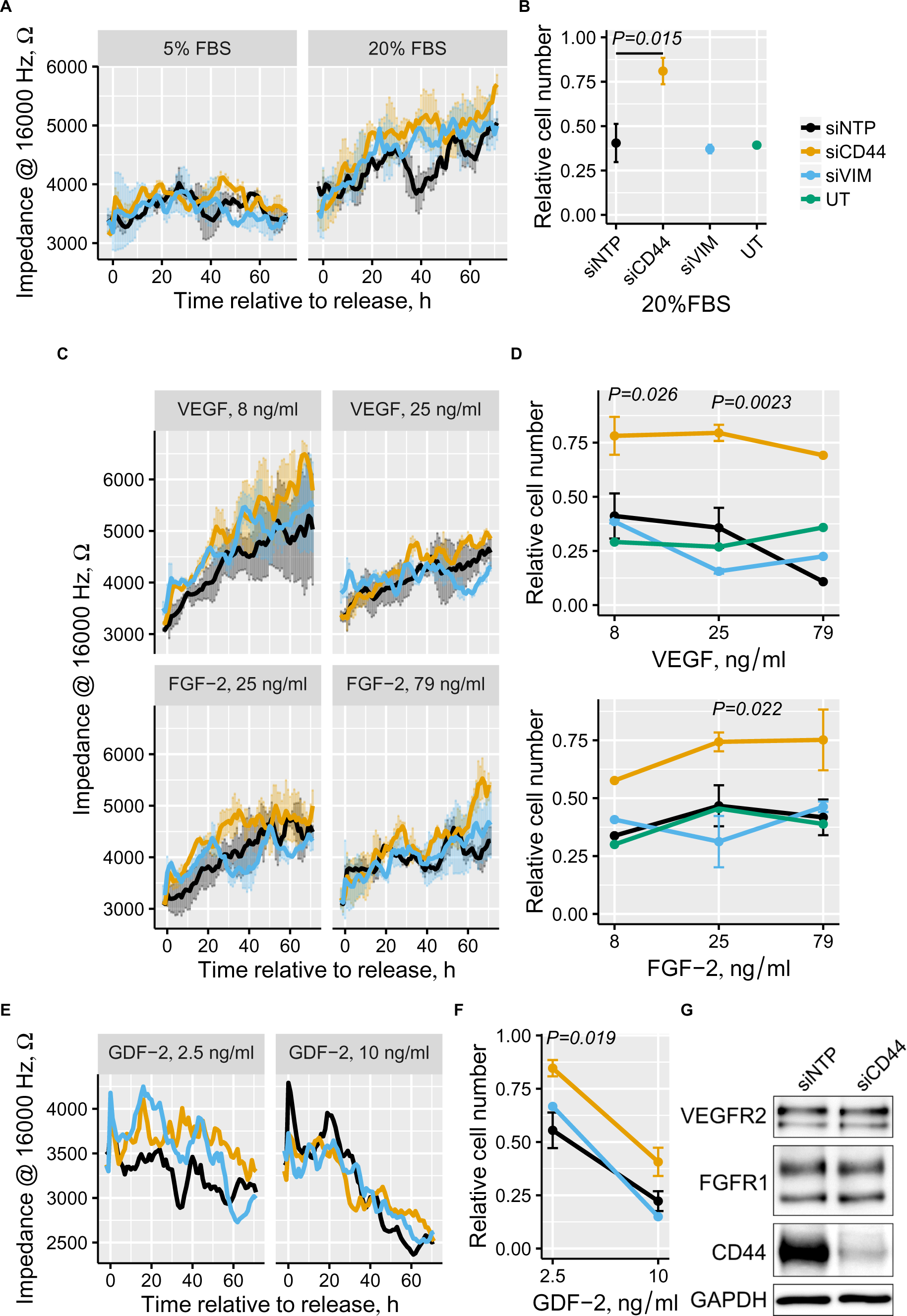
CD44 knockdown augments EC growth. siRNA transfected HUVECs were plated onto 96-well electrode arrays. After 24 h, the cells were starved in 1% FBS media overnight. After starving, the cells were released from cell cycle block by the addition of 20% FBS **(A)**; 8 ng/ml, 25 ng/ml, or 79 ng/ml FGF-2 or VEGF **(C)**; and 2.5 ng/ml or 10 ng/ml GDF-2 **(E)**. Following stimulation, HUVEC growth was monitored by recording electrode impedance. Raw impedance readings are shown to allow direct comparison to endpoint measurements. N = 2 for 5% and 20% FBS, 25 ng/ml and 79 ng/ml FGF-2, and 8 ng/ml and 25 ng/ml VEGF. **(B, D, F)** 72 h after release from cell cycle block the viable cell numbers were determined by measuring the ATP per well. Treatments are labeled as shown in (**B**). The data are represented as the mean ± SEM. P values are from the ANOVA post hoc comparisons using the Tukey HSD test. P values ≤ 0.05 of siCD44-siNTP comparisons are shown. siCD44-siNTP comparisons: N = 4 for 2.5 ng/ml and 10 ng/ml GDF-2, and N = 5 for 20% FBS, 25 ng/ml and 79 ng/ml FGF-2, and 8 ng/ml and 25 ng/ml VEGF. **(G)** Western blot analysis of CD44 silencing in HUVECs transfected with 30 nM siRNAs for 48 h. siNTP – non-targeting siRNA pool, siVIM – vimentin-targeting pool, siCD44 – CD44-targeting pool, UT – non-transfected cells.

Western blot analysis showed that FGFR1, VEGFR2 or activated VEGFR2 levels were not affected in CD44-silenced HUVECs (Fig. 5G and Supplemental Fig. 3C in Online Resource 1). The initial proliferation rate of CD44-silenced HUVECs after seeding and before serum deprivation was increased compared to non-targeting siRNA or untransfected controls, and CD44-silenced cells reached higher cell density within this time frame (Supplemental Fig. 6A in Online Resource 1). Notably, vimentin-silenced (siVIM) HUVECs showed similar behavior to CD44-silenced cells before serum starvation (Supplemental Fig. 6A in Online Resource 1). The observation that siCD44- or siVIM-transfected HUVECs reached higher cell density after seeding and before the start of serum starvation compared to siNTP-transfected or untransfected cells was confirmed by modeling impedance data for barrier formation (Supplemental Fig. 6B in Online Resource 1). The release of siCD44-transfected cells from serum starvation by VEGF or FGF-2 stimulation resulted in enhanced barrier reformation when compared to siNTP or siVIM-transfectants (Supplemental Fig. 6B in Online Resource 1).

We also assessed the effect of siCD44 by treating cells arrested in the cell cycle G1 phase with antimitogenic factor GDF-2. We found that GDF-2 induces cell cycle block and a subsequent decline in cell density, plausibly because cells undergo apoptosis (Fig. 4H). Impedance measurements showed that CD44-silencing did not rescue GDF-2-stimulated HUVECs from growth arrest and cell numbers declined over the course of the experiment (Fig. 5E).

However, the cell viability assay performed after impedance measurements showed that CD44-silencing resulted in more surviving cells compared to controls and partially rescued the cells from GDF-2-induced cell cycle block (Fig. 5F). Nevertheless, the GDF-2 dose-dependent inhibitory trend persisted.

Together, these experiments show that CD44 knockdown results in enhanced EC proliferation, irrespective of the specific growth factor used for stimulation. Furthermore, CD44-silencing experiments are consistent with *Cd44*^−/−^ mice data and suggest increased proliferation and survival of CD44-deficient ECs as a plausible cellular mechanism to enhance angiogenesis.

## Discussion

Here, we report that CD44 cell-surface glycoprotein is a negative regulator of angiogenesis. We show that CD44 constrains endothelial cell proliferation. Our results suggest that in the regulation of cell proliferation, CD44 functions independently of specific growth factor signaling pathways.

Based on our experiments, we extend the functions of CD44 to include the control of EC proliferation and angiogenesis. We found that blood vessel invasion into tumor extracellular matrix in response to FGF-2/VEGF stimulation was substantially increased in *Cd44*-null mice. This effect is likely to be cell-autonomous, as silencing of CD44 expression in cultured ECs also resulted in augmented cell proliferation. We suggest that CD44 functions downstream of mitogenic signaling. Griffioen et al. [28] have shown that CD44 is upregulated in response to FGF-2 or VEGF stimulation in cultured ECs, and in activated tumor blood vessels *in vivo*. Thus, enhanced angiogenesis and cell proliferation in case of CD44 deficiency or downregulation suggest that CD44 mediates negative feedback signaling that constrains cell proliferation. CD44 knockdown in dermal fibroblasts results in the stabilization of PDGF β-receptor and sustained ERK activation in response to PDGF-BB stimulation [29]. In our study, we show that intervening with CD44 function by silencing or CD44-3MUT-Fc has no effect on the activation of angiogenic growth factor receptors. However, we observed that the potency of CD44-3MUT-Fc to inhibit EC proliferation was inversely related to the potency of VEGF, FGF-2 or HGF to induce EC proliferation and survival. Several earlier reports have shown involvement of CD44 in TGF-β signaling [25–27]. Therefore, we tested if CD44 functions in GDF-2 signaling. We saw enhanced growth arrest and apoptosis of ECs in response to CD44-3MUT-Fc treatment in GDF-2-stimulated ECs. Nevertheless, our different *in vitro* experiments showed that GDF-2-mediated signaling is not affected by disrupting CD44 expression or increasing CD44 dose via CD44-3MUT-Fc. We conclude from these results that CD44 acts via a different mechanism than disrupting any specific growth factor pathways.

Plausibly, CD44-mediated negative feedback signaling on cell proliferation is activated by CD44-HA interaction. Binding of high molecular weight HA to CD44 controls proliferation of smooth muscle cells, and probably also other mesenchymal cell types, including ECs [10]. Kothapalli et al. [10] also showed that in *Cd44*-null mice the response to arterial injury resulted in increased neointima formation and smooth muscle cell proliferation during vessel regeneration. We have previously shown that the non-HA binding mutant of CD44 was as effective as its wild-type counterpart in angiogenesis inhibition in the chick chorioallantoic membrane angiogenesis model [11]. Here, we show that systemic administration of soluble mutant CD44 HABD (CD44-3MUT) has an antiangiogenic effect in a mouse model of angiogenesis, thus CD44-3MUT functions similarly to endogeneous CD44. In this context, we were interested whether endogeneous soluble CD44 levels correlate with angiogenesis. Soluble CD44 levels are reduced in immuno deficient BALB/c.Xid mice with defective B-cell maturation, and in SCID mice with absence of functional T cells and B cells, suggesting that immune cell-derived proteolytic activity is responsible for CD44 shedding [21]. In our angiogenesis assays, we observed that wild type mouse strains displayed much weaker angiogenic response than immuno deficient athymic nude mice. Given that nude mice carried normal *Cd44* gene dose, but their sCD44 levels were not known, we assumed that elevated angiogenesis in nude mice, compared to wild type strains, could be related to decreased sCD44 levels. We found serum sCD44 levels to be normal in athymic nude mice, suggesting that the induction of angiogenesis is not related to serum sCD44 levels. Furthermore, as athymic nude mice lack T cells, this result suggests that a large proportion of serum sCD44 is generated by B cell-dependent activity.

The signaling pathway downstream of CD44 is not well understood. CD44 has been implicated in cell–cell contact inhibition in schwannoma cells by recruiting the NF2 tumor suppressor protein to the plasma membrane [30]. Thus, it is possible that CD44 silencing abolishes the function of NF2, which leads to loss of contact inhibition and increased proliferation. However, embryonic fibroblasts isolated from *Cd44*-null mice still exhibit functional contact inhibition compared to cells from *Nf2*-null mice, but *Cd44*-null cells seem to display faster growth rates compared to wild-type cells [31]. Our impedance-based real-time monitoring of cell proliferation suggests steadily increased growth rates of CD44-silenced ECs after release from serum starvation. We found that barrier formation, which is directly related to cell density, is apparently more robust in CD44-silenced ECs after growth factor stimulation. This suggest that the mechanisms behind enhanced cell proliferation could be other than defective cell–cell adhesion.

Our *in vivo* findings contrast with previous works showing that CD44 absence or its downregulation *in vivo* results in reduced angiogenesis [3, 4]. Lennon et al. [4] studied the contribution of CD44 to HA oligomer-induced angiogenesis, and found that CD44 silencing *in vivo* resulted in inhibited angiogenesis in response to oligo HA. Cao et al. [3] used a relatively high number of rapidly growing B16 melanoma cells as a source of angiogenic growth factors in a Matrigel plug assay and allowed the blood vessels to grow for 5 days only. Nevertheless, tumor angiogenesis assays using two different cell lines with very different tumor growth kinetics, B16 melanoma and ID8-VEGF ovarian carcinoma, still suggested considerable inhibition of tumor formation and reduced vascular density at tumor margins in *Cd44*-null mice [3]. However, it is plausible that CD44-negative ECs were inhibited *in trans*by CD44 that was present on tumor cells [3]. Here, we show that administration of exogeneous soluble CD44 inhibited *in vivo* angiogenesis and EC proliferation. We found that *Cd44* heterozygous mice displayed angiogenesis at a similar level to wild-type animals, showing that *Cd44* is not haploinsufficient and lower than normal amounts are still sufficient for controlling angiogenesis. Tumor angiogenesis is dependent on interactions between tumor cells and host tissue stroma, and such interactions might be compromised in *Cd44*-null animals. Tumor cells recruit macrophages to promote angiogenesis. However, [3] showed that in case of *Cd44*-null mice bone marrow reconstitution with wild-type bone marrow did not rescue the angiogenesis defect, suggesting that endothelial CD44 expression is important.

*Cd44*-null mice develop normally and do not display apparent vascular abnormalities. We suggest that CD44 plays a non-redundant role in physiological angiogenesis. CD44-mediated interactions after its upregulation in endothelial cells in response to growth factor stimulation restrain cell proliferation. This control may contribute to the robust shutdown of angiogenesis during wound repair. In case of tumors, CD44-mediated control of angiogenesis might be overridden by a surplus of growth factors and increased shedding of CD44.

In summary, we conclude that CD44 functions as a negative regulator of angiogenesis. Therefore, systemic absence of CD44 expression in mice results in increased angiogenic response. Our results also demonstrate that soluble CD44 regulates angiogenesis by suppressing endothelial cell proliferation. Importantly, the antiangiogenic effect of CD44 is achieved independently of its HA-binding property. Together, our data suggest that CD44 is important in maintaining normal angiogenesis levels and targeting of CD44 can be utilized in antiangiogenesis treatment strategies for cancer or in other applications where angiogenesis modulation is desired.

## Materials and Methods

### Cells, Reagents and Primary Antibodies

HUVECs were obtained from Cell Applications, Inc., ECGS was from Millipore. VEGF-165 was from Serotec; GDF-2, HGF and FGF-2 were from Peprotech. Lipofectamine RNAiMAX (LF) was from Life Technologies. Non-targeting pool siRNA, #D-001810-10-05, human CD44 siRNA, #L-009999-00-0005, and human vimentin siRNA, #L-003551-00-0005 (ON-TARGETplus SMARTpool) were from Thermo Fisher Scientific. siRNA target sequences are listed in Supplemental Methods section in Online Resource 1. jetPEI-HUVEC transfection reagent was from Polyplus-transfection SA. Annexin V-FITC and annexin binding buffer were from BD Pharmigen. CellTiter-Glo reagent was from Promega. Primary antibodies, dilution and source used in this study: anti-CD44 (2C5) mouse mAb 1/1000 from R&D Systems, anti-VEGFR2 rabbit mAb (55B11) 1/1500 and anti-FGFR1 rabbit mAb (D8E4) from Cell Signaling Technology and anti-GAPDH mouse mAb 1/10000 from Millipore, IM7 rat anti-mouse CD44 (MCA4703; AbD Serotec), rat anti-mouse CD44 KM81-biotin (Abcam).

### Production of CD44-3MUT-Fc

CD44-3MUT with C-terminal human IgG1-Fc domain, recombinant human IgG1-Fc domain (rhIgG-Fc) and irrelevant human IgG1 mAb were produced by Icosagen Cell Factory (Estonia). Cystatin S signal peptide sequence was added to the N-terminus of the CD44-3MUT-Fc cDNA and the gene was synthesized by Genewiz, Inc. The synthesized CD44-3MUT-Fc cDNA was cloned into RSV-LTR promoter containing pQMCF-5 expression vector (Icosagen Cell Factory). The resulting expression plasmids were transfected into CHOEBNALT85 cells (Icosagen Cell Factory) and the expressed Fc-fusion proteins were purified by Protein G sepharose, followed by Superdex 200 gel-filtration chromatography. The purified CD44-3MUT-Fc had a monomeric molecular weight of approximately 60 kDa. The endotoxin level of the purified CD44-3MUT-Fc was < 10 EU/mg as determined by chromogenic Limulus amebocyte lysate test (Lonza).

### CD44-3MUT-Fc serum half-life

F344/NCrHsd male rats were from Harlan, Netherlands. The rats carried a polyurethane round tipped jugular vein catheter for blood sampling (Harlan Laboratories Surgical Services). After the pre-serum blood sample was taken, rats were injected intravenously via the tail vein with 3 mg of CD44-3MUT-Fc in 1 ml volume. Blood samples were collected using the jugular vein catheter at different time points. Blood samples were held at 37°C for 30 min to allow clot formation, and then centrifuged at 1300×g for 10 min at RT. The supernatants were collected and stored at −20°C until assayed. For sandwich ELISA microwell plates were coated with mouse anti-human IgG1 antibody clone G17-1 (BD Biosciences). Blocking was performed with 1.5% BSA/PBS. Standards were step-diluted (40 μg/ml – 0 μg/ml) in 0.5% BSA/PBS supplemented with 5%, 2% or 1% rat serum. Samples taken at different time points (pre-serum, from as soon as possible to 24 hours) were diluted 1:50 or 1:100 in 0.5% BSA/PBS solution and applied to wells. Biotin mouse anti-human IgG antibody clone G18-145 (BD Biosciences) and streptavidin-HRP were used for detection. Tetramethylbenzidine was used for color development. Concentration at time zero and half-life were estimated from two-parameter exponential decay model with the function *f*(*x*)=*d*(*exp*(−*x*/*e*)), where *d* is the upper limit at x = 0, and *e* is the decay constant.

### *In Vivo* Angiogenesis Assay

Animal experiments were conducted under the license of the Project Authorization Committee for Animal Experiments of the Ministry of Rural Affairs of the Republic of Estonia. We used the Directed in Vivo Angiogenesis Assay kit (DIVAA; Trevigen, USA) according to manufacturers instructions. For the angiogenesis assay, 20 μl angioreactors were filled with growth factor-reduced basement membrane extract containing 1.4 ng/μl FGF-2, 0.47 ng/μl VEGF and heparin for the induction of angiogenic response or an equal volume of PBS for uninduced controls. Angioreactors were implanted subcutaneously into the dorsolateral flank of 9-week-old athymic Nude-Foxn1/nu female mice (Harlan, Netherlands). *Cd44*-null mice were of mixed inbred background (B6;129-Cd44tm1Hbg/J), termed “Cd44KO mix”. B6;129 hybrid mice “WT mix” were used as controls. C57BL/6 and C3H mice were obtained from Harlan, Netherlands. The experimental groups of wild-type and *Cd44*-null mice comprised 8-11-week-old female and male mice. In one experiment, the Cd44KO mix and the respective wild-type mice were 40-43 weeks old. For the angiogenesis assay in the C57BL/6 genetic background, the *Cd44*-null allele was backcrossed for six generations to C57BL/6 mice (Harlan, Netherlands); experimental groups contained an equal number of female and male littermates. Implantation was performed on both flanks and one or two angioreactors were inserted per flank into immune competent mice or nude mice, respectively. Nude mice were injected intraperitoneally every second day with CD44-3MUT-Fc, irrelevant human IgG1 mAb, rhIgG-Fc or vehicle (PBS) during two weeks. Mice were sacrificed after 14 days from start of the experiment by carbon dioxide asphyxiation and angioreactors were dissected. The angioreactor contents were retrieved and the EC that had invaded the angioreactors were quantitated by FITC-Lectin (Griffonia simplicifolia lectin I) staining [32]. Cell-bound fluorescence was measured at 485 nm excitation and 535 nm emission wavelengths using a microtiter plate reader (Tecan, Swizerland). Data from irrelevant human IgG1 mAb and rhIgG1-Fc treatments were pooled for analysis. Raw fluorescence values from each experiment were scaled by dividing by their root mean square.

### Mouse sCD44 ELISA

For quantification of the serum levels of sCD44 in nude mice from the DIVAA experiments, wild-type mice of different backgrounds (C3H, C57BL/6, mixed), and their *Cd44*-null, heterozygous and wild-type littermates of the C57BL/6 background were used. For ELISA measurements 96-well plates were coated with 50 ng IM7 Ab/well overnight (ON) at 4°C and blocked with 5% non-fat dry milk for 2 h at 37°C. After blocking, samples and standards (Recombinant Mouse CD44 Fc Chimera; R&D Systems) were incubated in wells for 30 min at 37°C. Bound sCD44 was detected by incubating the plate with KM81-biotin Ab for 30 min at 37°C. For color development, Vectastain ABC (Vector Laboratories) and tetramethylbenzidine substrate were used. Absorbance was recorded at 450 nm using a microtiter plate reader (Tecan).

### HUVEC Growth and Treatments

HUVECs (passage 4-6) were cultured in 20% FBS containing M199 medium supplemented with 4 mM L-glutamine, 50 μg/ml heparin, 10 mM Hepes, and 30 μg/ml ECGS. For treatments, HUVECs were seeded into 0.1% gelatin-coated cell culture plates. After 24 h, the cells were starved ON in M199 media supplemented with 1% FBS, 25 mM Hepes and 4 mM L-glutamine. After starving, the cells were incubated with different concentrations of rhIgG-Fc or CD44-3MUT-Fc in the 5% FBS containing HUVEC growth media (5% FBS, M199, 4 mM L-glutamine, 12.5 μg/ml heparin, 10 mM Hepes, and 7.5 μg/ml ECGS) for 1 h at 37°C; and thereafter, stimulated with 25 ng/ml VEGF-165, 25 ng/ml FGF-2, 10 ng/ml GDF-2 or 63 ng/ml HGF. The cells were further grown for 48 or 72 h at 37°C.

### Electric Cell-Substrate Impedance Sensing Assay

Cell layer impedance was recorded by an ECIS Zθ instrument connected to a computer running an ECIS software version 1.2.169.0 (Applied Biophysics, USA). We used 96WE1+ PET plates, pretreated with 10 mM cysteine (Applied Biophysics, USA). HUVECs were seeded at a density of 5000 cells/well. The final media volume during treatments was 175 μl per well. Cell growth was monitored at seven frequencies in the range of 1000-64000 Hz. Each well was measured approximately four times per hour. To summarize different experiments, values for each experiment were binned by hours and hourly means were calculated. The release mark from serum starvation was set as time point zero. Hourly means of raw readings were normalized by dividing by the mean value of the first hour after time point zero.

### Apoptosis and Cell Viability Assays

For the apoptosis assay HUVECs were cultured in 0.1% gelatin-coated 24-well plates at a density of 25000 cells/well. Annexin V-FITC staining was performed according to manufacturer’s protocol. Briefly, cells were incubated with 1:20 Annexin V-FITC in 50 μl of annexin-binding buffer for 15 min at room temperature (RT) in the dark and analysed by FACS Calibur (BD Biosciences). For the cell viability assay, HUVECs were cultured in 0.1% gelatin-coated white 96-well cell culture plates (Greiner Bio-one) at a density of 5000 cells/well. After treatments, the cells were incubated with the CellTiter-Glo reagent (in a ratio of 1:1 of reagent volume to media) for 10 min at RT and luminescence was recorded using a microtiter plate reader. For analysis, raw luminescence values were min-max normalized. Curves were fitted using a five parameter logistic equation: $f(x) = Bottom + \frac{Top-Bottom}{(1+\exp(HillSlope(\log(x)-\log(EC50))))^S}$.

### HUVEC Transfection with siRNAs

siRNA transfections were performed using PEI or LF and 30 nM siRNAs. Transfection reagents and siRNAs were separately diluted in serum-free DMEM (4500 mg/l glucose). Diluted siRNA and transfection reagents were mixed and the transfection complex was incubated for 15-20 min at RT. The cells were transfected in 2% FBS and 4 mM L-glutamine containing DMEM (4500 mg/l glucose) for 3 h (PEI transfection) or 4 h (LF transfection) at 37°C. Then, the transfection media was replaced with 20% FBS HUVEC culture media and cells were further incubated for 24 h at 37°C. For impedance measurements, the transfected cells were plated out at a 5000 cells/well density into cysteine pretreated and gelatin coated 96WE1+ PET plates. After starving, cells were stimulated with different concentrations of GDF-2, VEGF or FGF-2 in 5% FBS containing HUVEC growth media. After 72 h of incubation, the CellTiterGlo reagent was added to cells for cell viability measurement.

### Western Blot Analysis

48 h after siRNA transfection, the cells were lysed in RIPA buffer (50 mM Tris pH 8.0, 150 mM NaCl, 1% NP-40, 0.5% sodium deoxycholate, 0.1% SDS) containing protease inhibitor cocktail (Roche). The proteins (5 μg of total protein) were separated on 7.5-10% SDS-PAGE gradient gel and transferred to PDVF membrane at 350 mA for 1.5 h. The membranes were blocked in 5% whey in 0.1% Tween20-TBS (TBST) at RT for 1 h, followed by a primary Ab incubation ON at 4°C and subsequent HRP-conjugated secondary Ab (Jackson Immunoresearch) incubation for 1 h at RT in 2% whey in TBST.

### Statistical Analysis

We used R version 3.3.1 (2016-06-21) for data analysis and graphs (for the complete list of packages see Supplemental Methods in Online Resource 1). The percentile confidence intervals were obtained using nonparametric bootstrap resampling on 1000 replications. Cohen’s d effect sizes with bootstrap confidence intervals were calculated using the *bootES* 1.2 package. Bayesian credible intervals were obtained via the Markov chain Monte Carlo method using the *rjags* 4.6 package. Scaling was done by dividing raw data by their root mean square using the R function *scale*. The data are shown as mean ± SEM.

### Electronic Supplementary Material

Supplemental Fig. 1 shows aortic fragment angiogenesis assay using aortic rings dissected from wild-type and *Cd44*^−/−^ mice. Supplemental Fig. 2 shows the CD44-3MUT-Fc serum half-life curve. Supplemental Fig. 3 shows the effect of CD44-3MUT-Fc on angiogenic growth factor receptor activation. Supplemental Fig. 4 shows the immunofluorescence analysis of GDF-2-induced pSMAD1/5 nuclear localisation in CD44-silenced HUVECs and transcription of SMAD target genes in CD44-silenced HUVECs in response to GDF-2 stimulation. Supplemental Fig. 5 shows the BMP-responsive element reporter activity of 10 ng/ml GDF-2 stimulated HUVECs transfected with CD44 siRNA and treated with CD44-3MUT-Fc and *in vivo* expression of SMAD target genes in mice treated with CD44-3MUT-Fc. Supplemental Fig. 6 shows that CD44 silencing augments EC growth and EC barrier formation is functional. Supplemental Table 1 gives the list of primers used for real-time qPCR experiments. Supplemental Table 2 gives the list of siRNA target sequences. Supplemental Table 3 gives the list of loaded R packages to compile this document.

## Author Contributions

Conceptualization, A.P., T.P., and A.V.; Methodology, A.P., M.S., and T.P.; Investigation, A.P., M.S., and T.P.; Formal Analysis, and Visualization, T.P.; Writing – Original Draft, A.P. and T.P.; Writing – Review & Editing, A.P., M.S., T.P., and A.V.; Funding Acquisition, A.V; Supervision, T.P., and A.V.

## Acknowledgments

We are grateful to Kati Mädo for her contribution to half-life studies and Aili Kallastu for her contribution to half-life and animal studies. We thank Richard Tamme and Alliki Lukk for proofreading the manuscript. This research was supported by the European Regional Development Fund via the Enterprise Estonia grants (EU28138/EU28658, EU30013) to the Competence Centre for Cancer Research and by the Estonian Science Fund Grant PUT698 to Andres Valkna. The authors declare no competing financial interests.

## Abbreviations List

CD44-3MUT: – CD44-HABD non-HA-binding triple-mutant
EC: – endothelial cell
ECIS: – Electric Cell-substrate Impedance Sensing
HA: – hyaluronan
HABD: – hyaluronan-binding domain
sCD44: – soluble CD44

## Compliance with Ethical Standards

Ethical Approval: All procedures performed in studies involving animals were in accordance with the ethical standards of the institution or practice at which the studies were conducted.

Conflict of Interest: The authors declare that they have no conflict of interest.

